# Reconstitution of CO_2_ regulation of SLAC1 anion channel and function of CO_2_-permeable PIP2;1 aquaporin as carbonic anhydrase 4 interactor

**DOI:** 10.1101/030296

**Authors:** Cun Wang, Honghong Hu, Xue Qin, Brian Zeise, Danyun Xu, Wouter-Jan Rappel, Walter F. Boron, Julian I. Schroeder

## Abstract

Daily dark periods cause an increase in the leaf CO_2_ concentration (*Ci*) and the continuing atmospheric [CO_2_] rise also increases *Ci*. Elevated *Ci* causes closing of stomatal pores thus regulating gas exchange of plants. The molecular signaling mechanisms leading to CO_2_-induced stomatal closure are only partially understood. Here we demonstrate that high intracellular 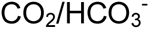 enhances currents mediated by the guard cell S-type anion channel SLAC1 when co-expressing either of the protein kinases OST1, CPK6 or CPK23 in *Xenopus* oocytes. Split-ubiquitin screening identified the PIP2;1 aquaporin as an interactor of the *β*CA4 carbonic anhydrase, which was confirmed in split luciferase, bimolecular fluorescence complementation and co-immunoprecipitation experiments. PIP2;1 exhibited CO_2_ permeability. Co-expression of *β*CA4 and PIP2;1 with OST1-SLAC1 or CPK6/23-SLAC1 enabled extracellular CO_2_ enhancement of SLAC1 anion channel activity. An inactive PIP2;1 point mutation was identified which abrogated water and CO_2_ permeability and extracellular CO_2_ regulation of SLAC1 activity in *Xenopus* oocytes. These findings identify the CO_2_-permeable PIP2;1 aquaporin as key interactor of carbonic anhydrases, show functional reconstitution of extracellular CO_2_ signaling to ion channel regulation and implicate SLAC1 as a bicarbonate-responsive protein in CO_2_ regulation of S-type anion channels.

## INTRODUCTION

Plant stomata, which are formed by pairs of guard cells in the epidermis of aerial tissues, control gas exchange and transpiration. Stomatal movements are regulated by several signals, including the phytohormone abscisic acid (ABA), CO_2_ (carbon dioxide), humidity, reactive oxygen species (ROS), light and pathogens (Hetherington and Woodward, 2003; Kim et al., 2010; Roelfsema et al., 2012; Murata et al., 2015). Daily respiration during darkness causes a rapid elevation in the CO_2_ concentration in the intercellular space in leaves (*Ci*) to ≥ 600 ppm (Hanstein et al., 2001). Furthermore, the continuing increase in the atmospheric CO_2_ concentration also causes an increase in *Ci*. Increased *Ci* reduces stomatal apertures and thus affects CO_2_ influx into plants, leaf heat stress, plant water use efficiency and yield (LaDeau and Clark, 2001; Medlyn et al., 2001; Hetherington and Woodward, 2003; Ainsworth and Long, 2005; Battisti and Naylor, 2009; Holden, 2009; Frommer, 2010).

Carbonic anhydrases catalyze the reversible reaction of 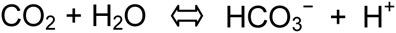. *Arabidopsis* double mutant plants in the *β*-carbonic anhydrases (*βCA1* and *βCA4*) display slowed stomatal movement responses to CO_2_ changes (Hu et al., 2010). Expression of an unrelated mammalian *α*-CA in *βca1βca4* double-mutant guard cells restores the stomatal CO_2_ response, suggesting that CA-mediated CO_2_ catalysis to bicarbonate and protons is an important step for transmission of the CO_2_ signal (Hu et al., 2010).

*SLAC1* (*SLOW ANION CHANNEL-ASSOCIATED 1*) encodes an S-type anion channel in *Arabidopsis* guard cells (Negi et al., 2008; Vahisalu et al., 2008). A group of protein kinases including OST1 (OPEN STOMATA 1), CPKs (Ca^2+^-dependent protein kinases) and GHR1 (GUARD CELL HYDROGEN PEROXIDE-RESISTANT 1) can phosphorylate and activate SLAC1 anion channels in *Xenopus* oocytes (Geiger et al., 2009; Lee et al., 2009; Geiger et al., 2010; Brandt et al., 2012; Hua et al., 2012). S-type anion channels are permeable to Cl^−^and 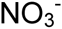, but not to 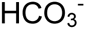 (Schmidt and Schroeder, 1994; Geiger et al., 2009; Xue et al., 2011). Intracellular bicarbonate generated by carbonic anhydrases acts as a second messenger and activates S-type anion channels in guard cells (Xue et al., 2011; Tian et al., 2015).

The OST1 protein kinase is required for CO_2_-induced stomatal closing (Xue et al., 2011; Merilo et al., 2013). A recent study reported that RHC1, a MATE-type transporter protein, acts as a bicarbonate sensor (Tian et al., 2015) and also inhibits HIGH LEAF TEMPERATURE 1 (HT1), a protein kinase that negatively regulates CO_2_-induced stomatal closing (Hashimoto et al., 2006). HT1 was found to phosphorylate and inhibit the OST1 protein kinase (Tian et al., 2015). Here, we have pursued investigation of the molecular targets and requirements for intracellular 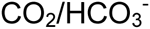 enhancement of SLAC1 anion channel activity.

We show that elevated intracellular NaHCO_3_ can enhance SLAC1 anion channel currents in both OST1-SLAC1 and CPK6/23-SLAC1 expressing oocytes. We isolate and characterize the PIP2;1 aquaporin as a new *β*CA4 carbonic anhydrase interactor and as a CO_2_-permeable aquaporin. In addition, we show that extracellular CO_2_ enhancement of S-type anion channels can be reconstituted when OST1-SLAC1 or CPK6/23-SLAC1 are co-expressed with the *β*CA4 carbonic anhydrase and PIP2;1 in *Xenopus* oocytes.

## RESULTS

### Elevated intracellular NaHCO_3_ can enhance SLAC1 anion channel currents in OST1-SLAC1 expressing oocytes

Intracellular bicarbonate enhances S-type anion channel currents in wild-type *Arabidopsis* guard cells (Hu et al., 2010; Xue et al., 2011; Tian et al., 2015). To test for minimal requirements by which bicarbonate could regulate the S-type anion channel SLAC1, we expressed SLAC1yc, or co-expressed SLAC1yc with the OST1yn protein kinase in *Xenopus* oocytes. To investigate electrophysiological responses, we injected 11.5 mM NaHCO_3_ into oocytes, mimicking high bicarbonate conditions that activate S-type anion channel currents in guard cells (Hu et al., 2010; Xue et al., 2011). Injection of 11.5 mM NaHCO_3_ buffered to pH 7.5 with Mes/Tris corresponds to 10.5 mM free 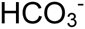 and 1 mM free CO_2_ (see Methods). Results from over six independent oocyte batches showed that high NaHCO_3_ consistently enhanced SLAC1yc/OST1yn-mediated anion channel currents in oocytes (Figure 1; p = 0.027 at -160 mV in SLAC1yc/OST1yn vs SLAC1yc/OST1yn + 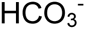). In contrast, SLAC1yc expression alone or water-injected control oocytes showed no significant NaHCO_3_ activation (Figure 1). To determine whether this activation is pH dependent, we also injected 11.5 mM NaHCO_3_ buffered to pH 7 and pH 8 into oocytes. The results showed that pH had no effect on the activation of SLAC1 anion channel currents (Supplemental Figure 1). In additional experiments, we injected an iso-osmotic 23 mM sorbitol solution buffered to pH 8. No enhancement of SLAC1-mediated ionic currents was observed, suggesting that a pH change was not the mechanism mediating enhancement of SLAC1-mediated currents. As an additional control for NaHCO_3_, injection of NaCl at the same concentration into oocytes did not enhance SLAC1yc/OST1yn-mediated anion channel currents, but rather showed an average reduction in anion currents (Supplemental Figure 2).

**Figure 1.**
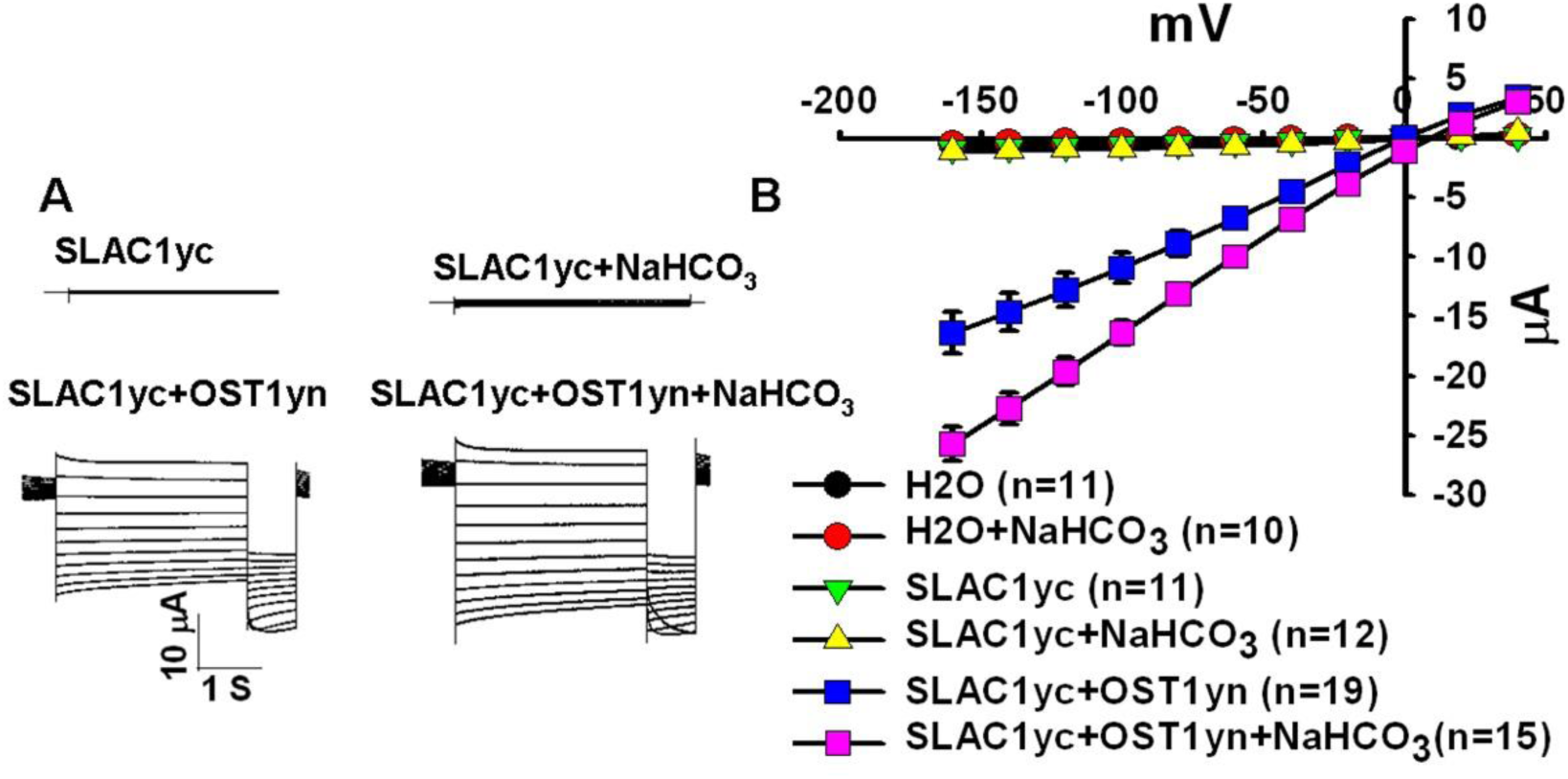
Injection of 11.5 mM NaHCO_3_ into oocytes causes enhancement of SLAC1-mediated anion channel currents when SLAC1 is co-expressed with OST1, while SLAC1 expression alone or water-injected oocytes did not show significant 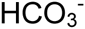 regulation. **(A)** Whole-cell currents were recorded from oocytes expressing the indicated cRNAs. During recordings of SLAC1 anion currents, single voltage pulses were applied in −20 mV decrements from +40 to −160 mV for 4 s and the holding potential was 0 mV. **(B)** Steady-state current-voltage relationships from oocytes recorded as in (A). Numbers of oocytes recorded for each condition in the same batch of oocytes are provided in the current-voltage panels in all figures. Data are mean ± s.e.m. For some data points the symbols were larger than the s.e.m. and therefore error bars are not visible in current-voltage plots in (B). Results from over five independent batches of oocytes showed similar results.

We investigated whether the 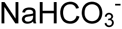 induced enhancement of SLAC1yc/OST1yn-mediated anion channel currents was dependent on the NaHCO_3_ concentration. A series of final intracellular NaHCO_3_ concentrations was microinjected: 0 mM, 1 mM, 5.7 mM and 11.5 mM. Results from over three independent oocyte batch experiments showed that low NaHCO_3_ failed to significantly enhance SLAC1yc/OST1yn-mediated anion channel currents, while injection of high NaHCO_3_ concentrations (5.7 mM and 11.5 mM) enhanced SLAC1yc/OST1yn-mediated anion channel currents (Figure 2A and 2B), which is in line with required NaHCO_3_ concentrations in guard cells (Xue et al., 2011., Tian et al., 2015). Furthermore, activation by 11.5 mM NaHCO_3_ was stronger than 5.7 mM NaHCO_3_ (Figure 2A and B; p = 0.034 at -160 mV in 11.5 mM NaHCO_3_ vs 5.7 mM NaHCO_3_). Thus high intracellular NaHCO_3_ concentrations could enhance SLAC1yc/OST1yn-mediated anion channel currents at similar concentrations as in guard cells (Xue et al., 2011).

**Figure 2.**
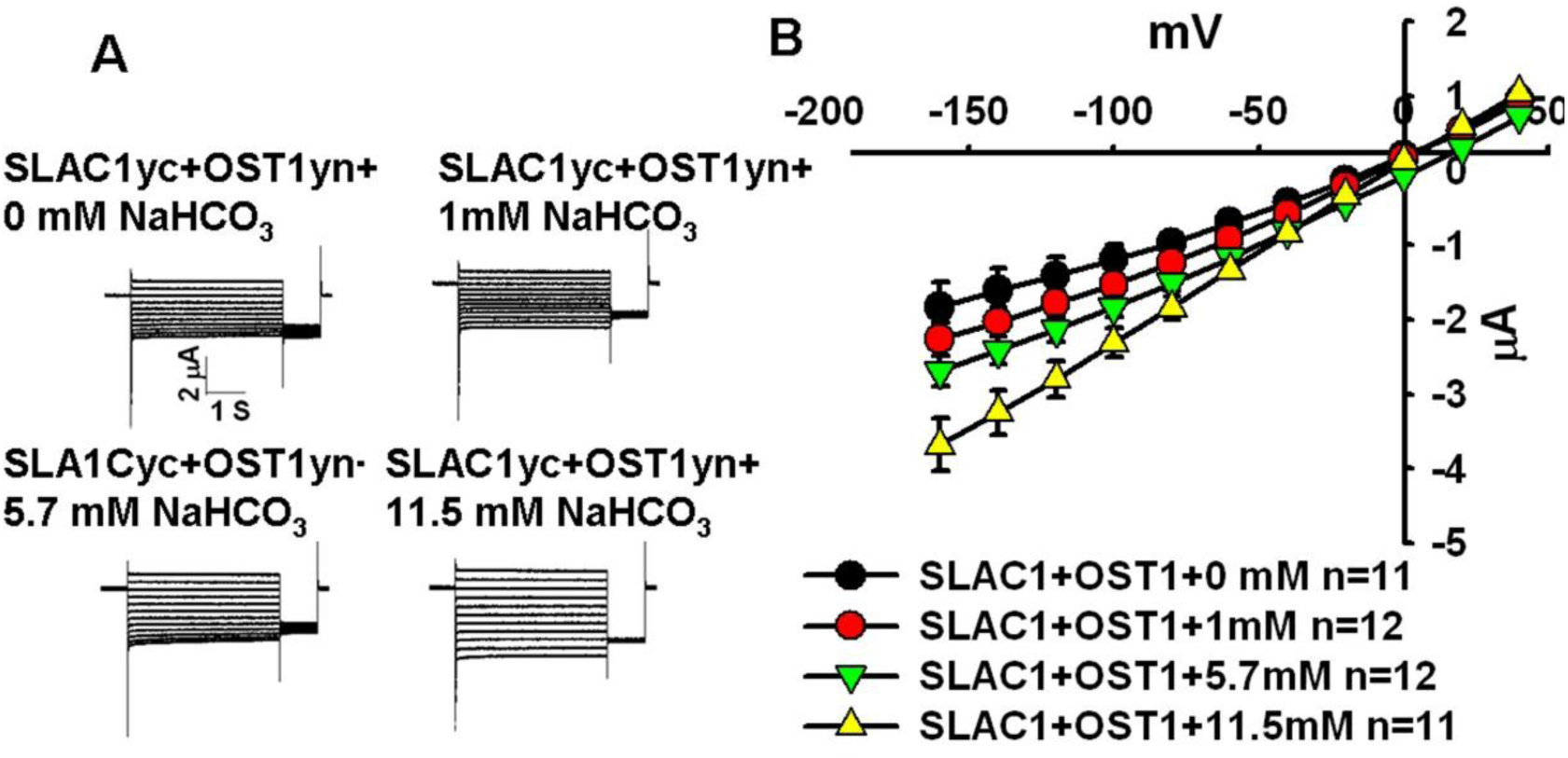
NaHCO_3_ concentration-dependent enhancement of SLAC1-mediated anion channel currents when SLAC1 was co-expressed with OST1 in *Xenopus* oocytes. **(A)** Whole-cell currents were recorded from oocytes after injection of the indicated final concentrations of NaHCO_3_. **(B)** Steady-state current-voltage relationships from oocytes recorded as in (A). Data are mean ± s.e.m. For some data points the symbols were larger than the s.e.m. and therefore the error bars are not visible for some data points in (B). Results from three independent oocyte batches showed similar results.

### PIP2;1 aquaporin as a new βCA4 carbonic anhydrase interactor and as a CO_2_-permeable aquaporin

The *β*CA1 and *β*CA4 carbonic anhydrases function in CO_2_-induced stomatal closing (Hu et al., 2010; Hu et al., 2015). To characterize the CO_2_ signaling mechanisms mediated by carbonic anhydrase proteins, we screened for interactors of these *β*-carbonic anhydrases by using the yeast two hybrid system with an *Arabidopsis* cDNA library (BD Clontech), using full length *β*CA4 cDNA as bait. However, no reproducible candidate interactors were isolated. The *β*CA4 carbonic anhydrase has been reported to be targeted to the plasma membrane in transiently-transformed tobacco cells, and *β*CA1 was mainly targeted to chloroplasts (Fabre et al., 2007; Hu et al., 2015). To screen for putative plasma membrane interactors of the carbonic anhydrase *β*CA4, the split-ubiquitin system (SUS) was developed and improved to detect protein-protein interactions (Jones et al., 2014). We used *β*CA4 as a bait and screened a SUS cDNA library, which was constructed in the pNX33-DEST vector (Grefen et al., 2007). In this screen, reproducible putative *β*CA4-interacting proteins were isolated, including Nodulin MtN3 (At3g06433), CNGC13 (At3g01010) and the PIP2;1 aquaporin (Figure 3A).

**Figure 3.**
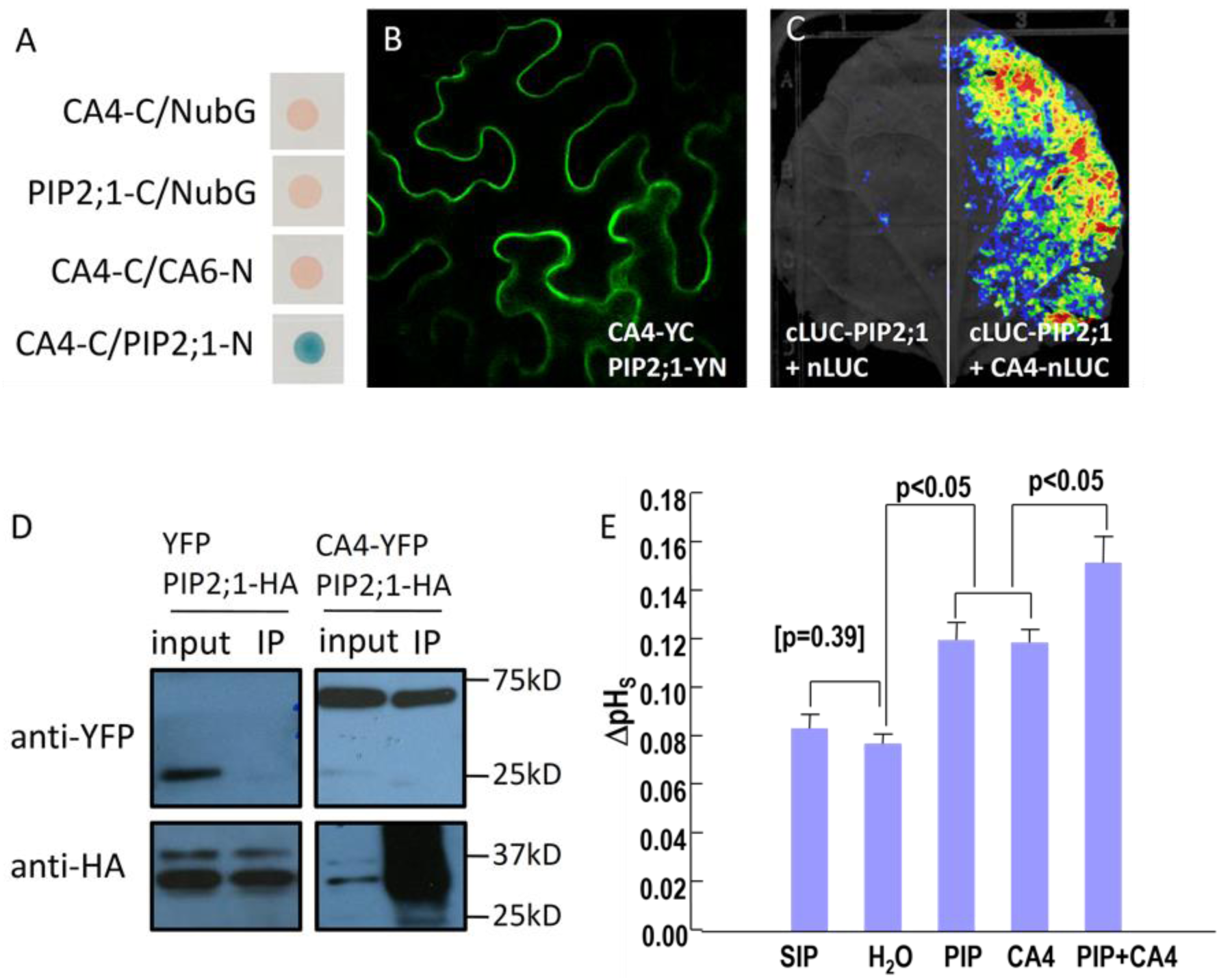
The carbonic anhydrase *β*CA4 interacts with the PIP2;1 protein and *β*CA4 increases the CO_2_ permeability mediated by PIP2;1. **(A)** The aquaporin PIP2;1 was identified as a *β*CA4-interacting protein by screening a split ubiquitin yeast two hybrid library. Interactions were shown in blue on the medium plus X-gal substrate. *β*CA6 and soluble NubG were used as negative controls. **(B, C)** Split YFP (B) and reversible split luciferase complementation assays (C) showed that *β*CA4 interacts with PIP2;1 at the plasma membrane in tobacco leaves. Note that reversible split luciferase assays do not permit single cell resolution and a whole leaf is shown in (C). **(C, left)** cLUC-PIP2;1 with only nLUC was used as negative control, and showed no clear luciferase bioluminescence signal. **(D)** Co-immunoprecipitation experiments show an interaction of *β*CA4 with the PIP2;1 aquaporin. Crude protein extracts from inoculated *Nicotiana benthamiana* leaves were used for immunoprecipitation as an input. Input protein was immuno-precipitated with anti-HA matrix. The input and immuno-precipitate (IP) were probed with anti-HA or anti-GFP antibodies as indicated. **(E)** *β*CA4 and PIP2;1 enhance the CO_2_ permeability of *Xenopus* oocytes as analyzed by membrane surface changes in pH. SIP1A-(At3g04090) & H_2_O-injected oocytes serve as control injections (* P < 0.05).

To further investigate putative interactors of *β*CA4 and assess whether these interactions occur *in vivo*, PIP2;1 aquaporin was chosen for further analyses in the present study. The split YFP combinations of *β*CA4-YC with PIP2;1-YN were transiently co-expressed in *Nicotiana benthamiana* leaves. Bimolecular fluorescence (BIFC) (Walter et al., 2004; Bracha-Drori et al., 2004) results showed that *β*CA4-YC and PIP2;1-YN interacted with each other in the vicinity of the plasma membrane (Figure 3B), supporting the SUS results. Reversible split luciferase complementation assays (Chen et al., 2008) were conducted to further test protein-protein interactions in *Nicotiana benthamiana* leaves and also showed that *β*CA4 interacts with PIP2;1 using this approach in leaves (Figure 3C). Furthermore, co-immunoprecipitation experiments were performed to test for protein-protein interactions in *Nicotiana benthamiana* leaves. *β*CA4-YFP co-immunoprecipitated with PIP2;1-HA as detected using HA and GFP antibodies (Figure 3D). In summary, four independent approaches: SUS, BiFC, split luciferase complementation and co-immunoprecipitation showed interactions of *β*CA4 and PIP2;1 in *vitro* and in plant cells.

The aquaporins NtAQP1 from tobacco, PIP1;2 from *Arabidopsis* and four PIP2 proteins in barley, were identified as CO_2_ transporters in plant cells (Uehlein et al., 2003; Uehlein et al., 2008; Uehlein et al., 2012; Mori et al., 2014). We investigated whether PIP2;1 can also mediate CO_2_ transport. PIP2;1 was expressed in *Xenopus* oocytes. pH changes in the proximity of the plasma membrane were used as an indicator that reflect the changes in acidification at the membrane surface of *Xenopus* oocytes mediated by CO_2_ flux (Musa-Aziz et al., 2009, 2014). The data showed that expression of PIP2;1 resulted in an enhanced change in the surface pH of oocytes, indicating that PIP2;1 enhanced the CO_2_ permeability of oocytes over the background CO_2_ permeability (Figure 3E and Supplemental Figure 3). Interestingly, when PIP2;1 was co-expressed with *β*CA4 in *Xenopus* oocytes, ΔpH was significantly increased compared to either PIP2;1 or *β*CA4 expression alone, suggesting enhanced CO_2_ transport into oocytes (Figure 3E and Supplemental Figure 3), which is similar to human carbonic anhydrasell-enhanced CO_2_ fluxes across *Xenopus* oocytes plasma membranes (Musa-Aziz et al., 2014).

### Reconstitution of extracellular CO_2_ signaling to SLAC1 anion channel regulation

As *β*CA4 physically interacts with the PIP2;1 aquaporin (Figure 3A-D) and PIP2;1 shows a CO_2_ permeability (Figure 3E), we pursued experiments to investigate whether extracellular 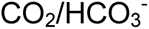 regulation of anion channels can be reconstituted in *Xenopus* oocytes.

Although injection of NaHCO_3_ into oocytes enhanced SLAC1-mediated currents (Figures 1 and 2), increasing extracellular 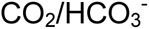 by addition of 11.5 mM NaHCO_3_ to the bath solution did not enhance SLAC1yc/OST1yn expressing oocyte currents (Figure 4A). When either *β*CA4 or the PIP2;1 aquaporin alone were co-expressed with SLAC1yc and OST1yn in *Xenopus* oocytes in the presence of high 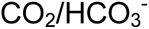 in the bath solution, no significant increase in ion currents was observed in three independent batches of oocytes (Figure 4B). We then co-expressed *β*CA4, PIP2;1 with SLAC1yc and OST1yn in oocytes, and found *β*CA4 and PIP2;1 did not significantly enhance ion currents in the bath solution without extracellular addition of NaHCO_3_ (Figure 4C). In contrast, in more than four independent batches of oocytes in the presence of extracellular NaHCO_3_, co-expression of *β*CA4, PIP2;1, SLAC1yc and OST1yn in oocytes displayed larger anion channel currents than SLAC1yc/OST1yn co-expressing oocytes (Figure 4D and 4E, p = 0.017 at -160 mV in SLAC1yc/OST1yn vs SLAC1yc/OST1yn+CA4+PIP2;1).

**Figure 4.**
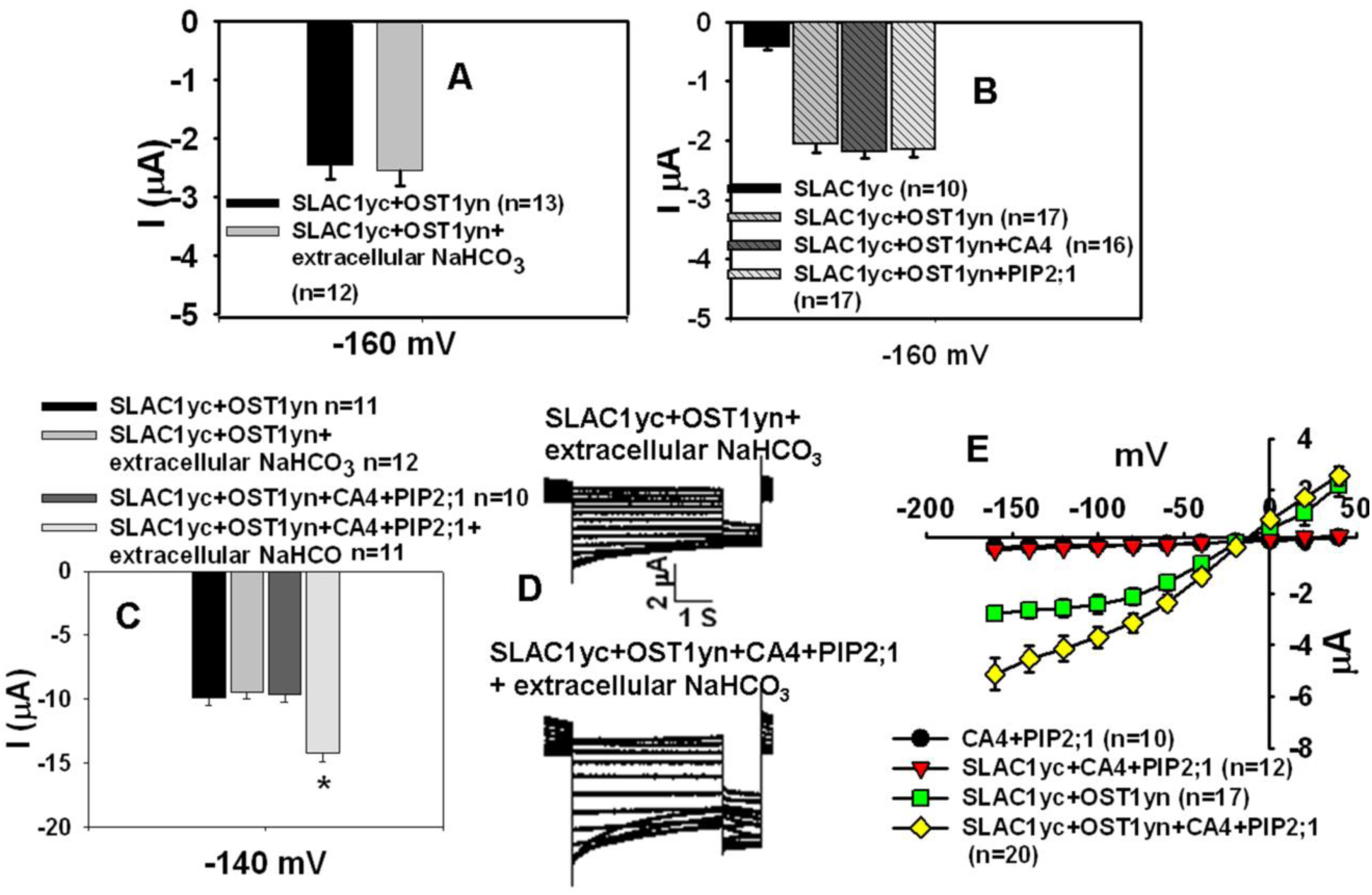
Reconstitution of extracellular 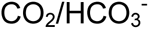enhancement of SLAC1-mediated anion channels requires PIP2;1 and *β*CA4 in *Xenopus* oocytes. **(A)** Increasing extracellular 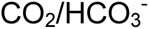by addition of 11.5 mM NaHCO_3_ did not enhance in SLAC1yc/OST1yn-expressing oocytes. **(B)** Either *β*CA4 or PIP2;1 expressed alone together with SLAC1 and OST1 did not suffice to enhance SLAC1-mediated anion channel currents in *Xenopus* oocytes. Data in (A) and (B) are shown at −160 mV as mean ± s.e.m. **(C)** Co-expression of *β*CA4 and PIP2;1 with SLAC1yc/OST1yn did not significantly enhance ion currents in the bath solution without high 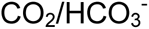. **(D)** Whole cell currents were recorded from oocytes co-expressing the indicated cRNAs. *β*CA4 and PIP2;1 co-expression enhanced SLAC1/OST1-mediated anion channel currents in the presence of 11.5 mM NaHCO_3_ in the bath solution. **(E)** Steady-state current-voltage relationships from oocytes recorded as in (D). Data are mean ± s.e.m. The results were found in at least three independent batches of oocytes.

### Functional PIP2;1 is required for the extracellular CO_2_ response

To test the hypothesis that a functional PIP2;1 is required for the extracellular CO_2_ response, we attempted to design non-permeable PIP2;1 isoforms. Several aquaporin structures have been resolved by X-ray crystallography, including human hAQP1, human hAQP5 and spinach aquaporin soPIP2 (Ren et al., 2000; Fujiyoshi et al., 2002; Kukulski et al., 2005; Horsefield et al., 2008; Nyblom et al., 2009). We then aligned hAQP1, hAQP5, SoPIP2 with PIP2;1. Based on this model, we speculated that L81; W85 and F210 might play a role in CO_2_ permeability. These three PIP2;1 amino acid residues were mutated to alanine in PIP2;1, and the cRNAs were expressed in *Xenopus* oocytes. In addition, grapevine VvPIP2;5 expressing oocytes have a very low water permeability compared with VvPIP2;1 and SoPIP2;1 (Shelden et al., 2009). Previous sequence alignments of the conserved B loop led to the model that W100 is a large and hydrophobic residue that may block the pore of VvPIP2;5 (Shelden et al., 2009). We therefore mutated the corresponding residue in PIP2;1-G103 to W. The osmotic water permeability (Pf) of PIP2;1-L81A, PIP2;1-W85A and PIP2;1-F210A showed no significant difference compared to the water permeability of wild type PIP2;1 (Figure 5A). Interestingly, the PIP2;1-G103W point mutant isoform exhibited a much weaker water permeability than wild type PIP2;1 in *Xenopus* oocytes (Figure 5A). This supports the model that the PIP2;1-G103 residue is important for PIP2;1-mediated function.

**Figure 5.**
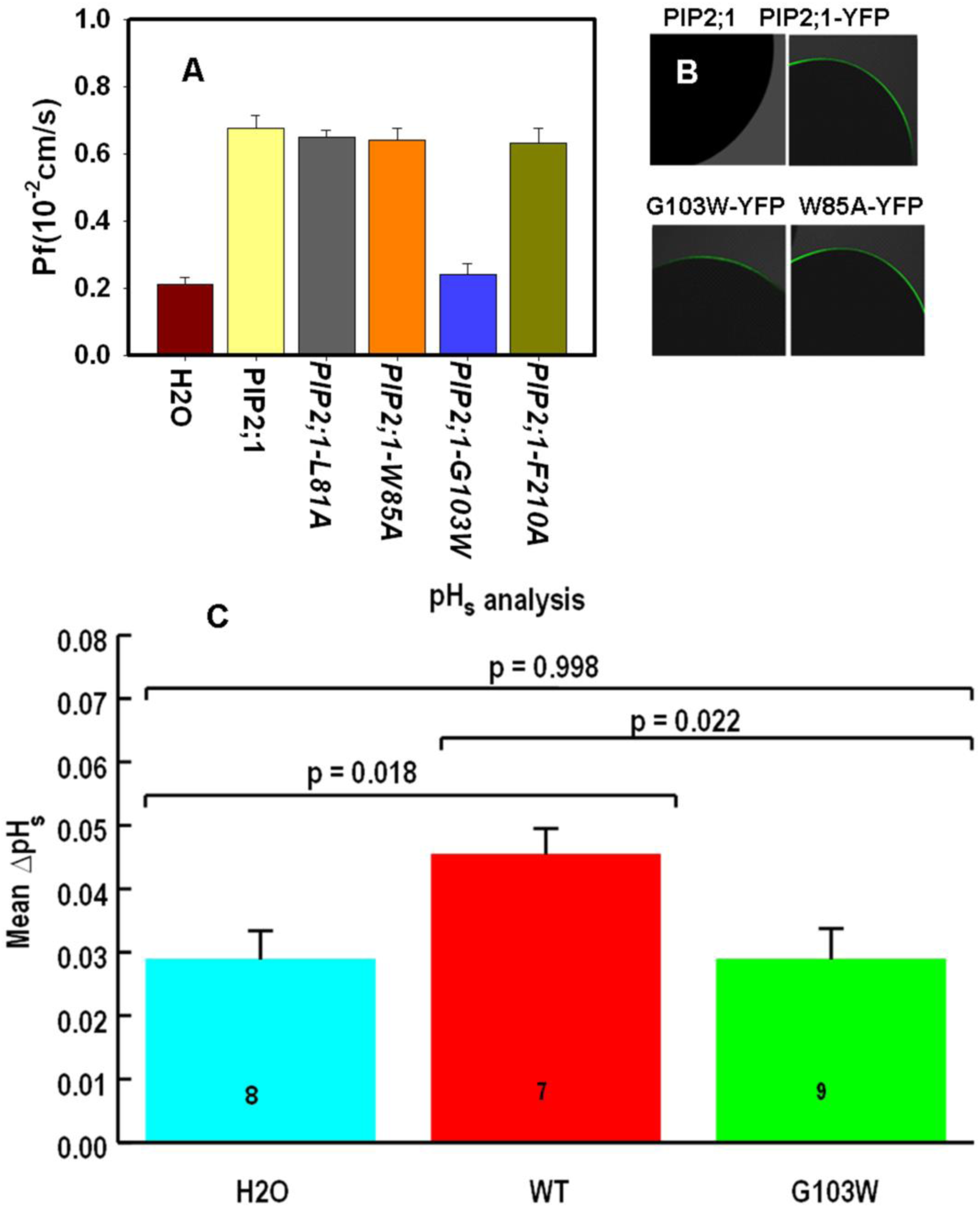
Osmotic water permeability coefficient, PIP2;1 location in *Xenopus* oocytes and surface pH analyses. **(A)** *PIP2;1-G103W-* and non-PIP2;1-expressing control oocytes showed a low water permeability. Results are shown as means ± s.e.m measurements from five to eight oocytes from one batch of oocytes. **(B)** PIP2;1 and its point mutation isoforms as YFP-fusions were present in the vicinity of the oocytes plasma membrane, PIP2;1 alone served as a negative control. **(C)** *PIP2;1-G103W* impaired the CO_2_ permeability of PIP2;1 as measured by changes in pH_s_.

To determine whether the PIP2;1 mutant protein isoforms are expressed and translocated to the plasma membrane of *Xenopus* oocytes, mRNAs of PIP2;1-YFP fusion proteins were expressed in oocytes. Confocal fluorescence microscopy analyses showed that all tested PIP2;1 mutant isoforms, including the non-functional PIP2;1-G103W-YFP fusion protein, were present at the plasma membrane (Figure 5B).

We next co-expressed the four mutant PIP2;1 isoforms with *β*CA4, SLAC1yc and OST1yn in *Xenopus* oocytes. We found that expression of the mutants PIP2;1-L81A, PIP2;1-W85A and PIP2;1-F210A could still mediate extracellular 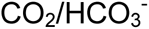 enhancement of SLAC1yc/OST1yn-mediated anion channel currents in four independent oocyte batches (Figure 6 and Supplemental Figure 4). In contrast the PIP2;1-G103W mutant isoforms did not enable extracellular 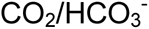-mediated enhancement SLAC1yc/OST1yn-mediated anion channel currents in three independent oocytes batches (Figure 6 and Supplemental Figure 4).

**Figure 6.**
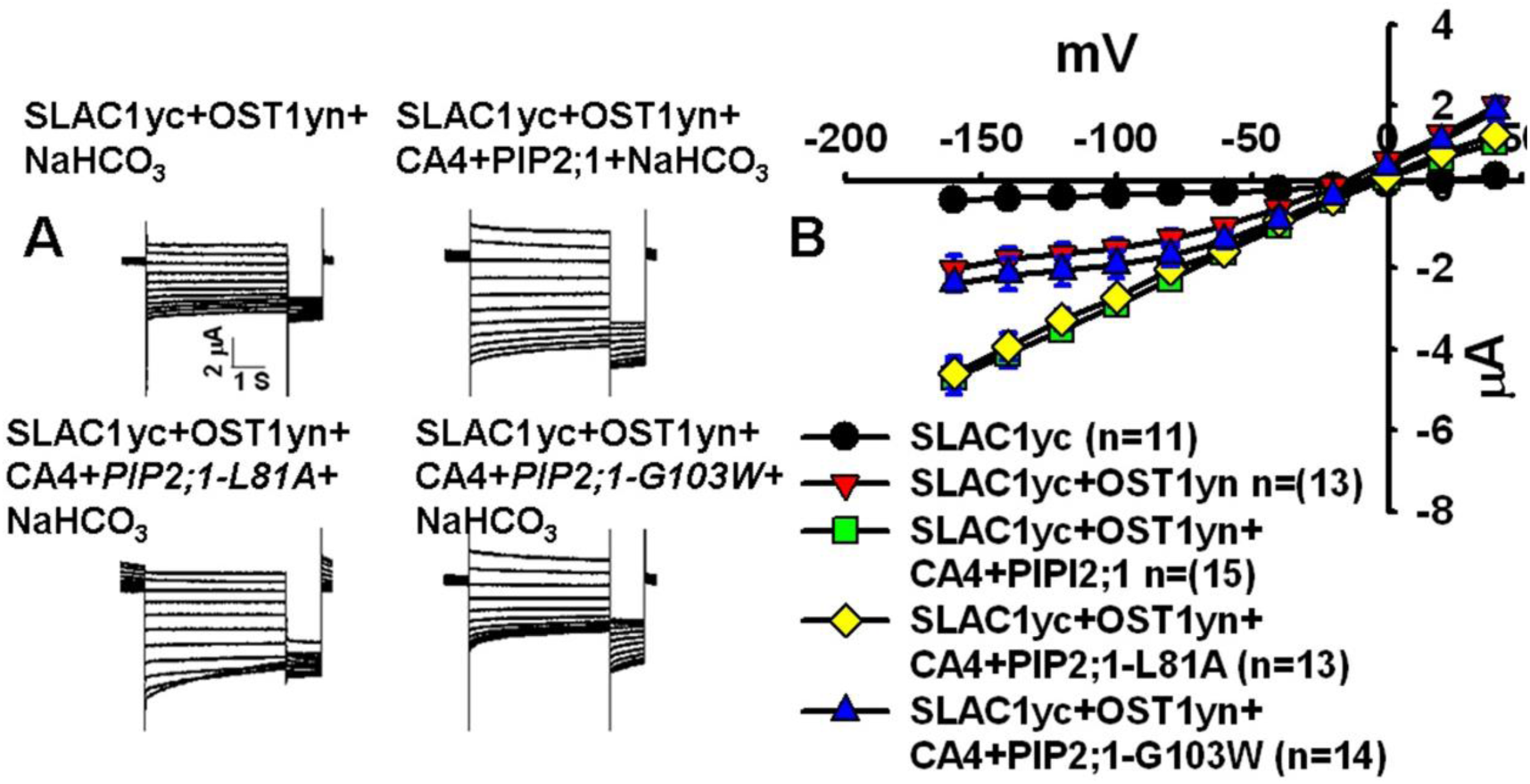
The *PIP2;1-G103W* mutant disrupted extracellular 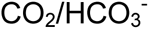-induced PIP2;1-*β*CA4 enhancement of SLAC1/OST1-mediated anion channel currents in oocytes. **(A)** Whole-cell currents were recorded from oocytes expressing the indicated cRNAs with 11.5 mM NaHCO_3_ in the bath solution. The voltage protocol was the same as in Figure 1. **(B)** Steady-state current-voltage relationships from oocytes recorded as in (A). Data are mean ± s.e.m. Results from three independent experiments showed similar results.

The PIP2;1-G103W mutation disrupted water permeability but not plasma membrane localization of PIP2;1 in *Xenopus* oocytes (Figure 5B). We investigated whether PIP2;1-G103W affects CO_2_ transport in oocytes. Experiments showed that expression of PIP2;1 resulted in an enhanced change in surface pH of oocytes, whereas in the same oocytes batches PIP2;1-G103W did not significantly changed the surface pH compared to control oocytes (Figure 5C). These results suggested that the G103W mutation impaired both PIP2;1-mediated water and CO_2_ transport (see discussion).

Surface pH changes of oocytes suggested that PIP2;1 and *β*CA4 co-expression together enhanced CO_2_ transport into oocytes more than expression of either protein alone (Figure 3E). We pursued mathematical modeling (Somersalo et al., 2012) to simulate PIP2;1 as enhancing the oocyte plasma membrane CO_2_ permeability and *β*CA4 which enhances CO_2_ catalysis in oocytes. We modeled the response of an oocyte following a sudden increase in external CO_2_ concentration (see methods). Simulations were started with the internal CO_2_ concentration set to 200 ppm and the external concentration increased from 200 ppm to 800 ppm (Hu et al., 2015). This sudden jump results in an influx of CO_2_ and a transient increase in the membrane surface pH, pH_s_ (Supplemental Figure 5).

Simulations show the predicted pH_s_ as a function of time for an oocyte for our baseline parameters (i.e., low membrane permeability and no carbonic anhydrase mediated acceleration of CO_2_ catalysis) as a black line (Supplemental Figure 5). Consistent with Somersalo et al (2012), the predicted pH_s_ increases, reached a transient maximum. The effect of carbonic anhydrases was simulated by setting an acceleration factor for CO_2_ catalysis of F to 100 in the interior of oocytes. As a result of the accelerated dynamics, the predicted CO_2_ influx was increased, resulting in an increase in the pH_s_ values (Supplemental Figure 5). We also simulated the effect of PIP2;1 expression, without the presence of CA, by increasing the membrane permeability to P_M,CO2_=10 cm/s. As expected, this increase in membrane permeability increased the predicted CO_2_ influx and the peak value of pH_s_ (green line). Finally, we modeled the combined effects of PIP2;1 and carbonic anhydrases co-expression by setting P_M,CO2_=10 cm/s and F=100. Consistent with the experiments, this combination led to an even higher predicted increase in value of ΔpH_s_ (blue line). Thus this model for CO_2_ dynamics can capture the relative larger increase in ΔpH_s_ upon co-expression of PIP2;1 and CA. Together with the experimental results (Figure 3E and Supplemental Figure 3), a threshold CO_2_ influx may be required to produce a measurable change in SLAC1 activity.

### PIP2;1 mutation alone did not significantly impair CO_2_- and ABA-regulation of stomatal movements

To explore whether *PIP2;1 insertional* mutation in the PIP2;1 gene in *Arabidopsis* alone is sufficient to impair CO_2_-regulation of stomatal movements, we pursued stomatal conductance analyses in response to CO_2_ concentration changes in *PIP2;1* T-DNA mutant plants. However, *pip2;1* mutant leaves showed similar stomatal responses to CO_2_ changes as wildtype leaves (Figure 7A and 7B).

**Figure 7.**
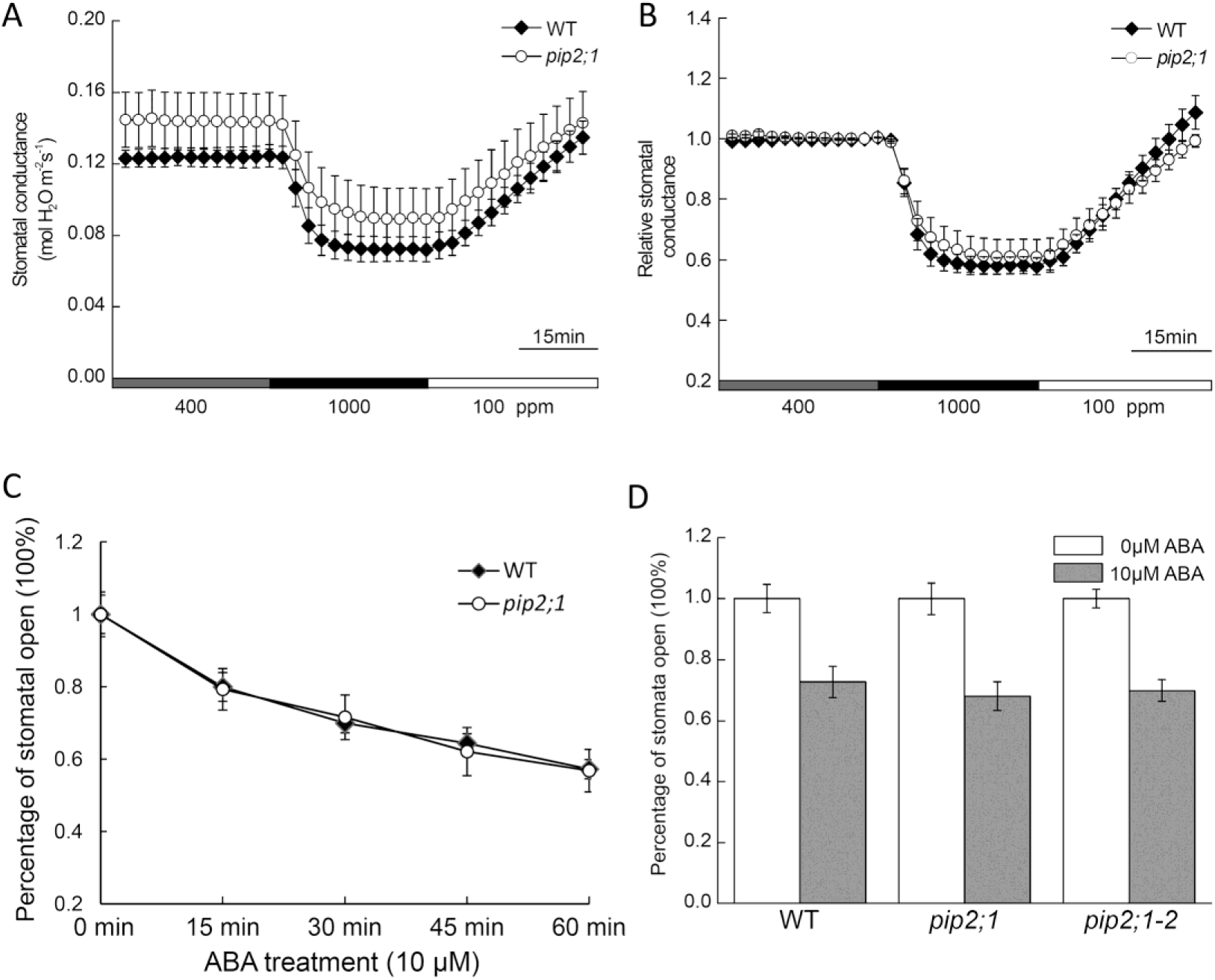
*PIP2;1* mutation alone did not significantly impair CO_2_-and ABA-regulation of stomatal movements. **(A)** Time-resolved intact leaf stomatal conductance in *pip2;1* (CS320492) and wildtype Col (WT) plants with [CO_2_] shifts indicated at the bottom. **(B)** Relative stomatal conductance data shown in (A). **(C)** Time-coursed analysis of stomatal movements in response to ABA treatment in individual mapped stomata in wildtype and *pip2;1* mutant. For these analyses individual stomata were imaged and tracked (Xue et al., 2011). **(D)** Stomatal ABA responses were analyzed in two different *pip2;1* alleles. n = 3 experiments, 30 stomata per experiment and condition, genotype blind. Data are mean ± s.e.m.

A recent study reported that *PIP2;1* knockout mutant plants have a defect in stomatal closure, specifically in response to ABA (Grondin et al., 2015). We also investigated whether PIP2;1 is involved in the ABA-induced stomatal closing pathway, since the same aquaporins are known to transport water and CO_2_ (Mori et al., 2014). Genotype-blind stomatal movement imaging analyses of individually mapped stomata showed that *pip2;1* single mutant stomatal retained intact responses to 10 μΜ ABA treatment in time-course analyses of ABA-induced stomatal closing (Figure 7E). Furthermore, we also tested the effect of ABA on another published *pip2;1* mutant *pip2;1-2* (Grondin et al., 2015) in parallel, but stomata in both *pip2;1* and *pip2;1-2* mutant leaf epidermal layers closed to similar levels as wildtype one hour after ABA treatment (Figure 7F). These data indicated that PIP2;1 mutation alone was insufficient under the imposed conditions to impair the ABA-induced stomatal closing pathway. These relevant findings may be related to co-expression of multiple *PIP* homologs in *Arabidopsis* guard cells (Yang et al., 2008; Zhao et al., 2008; Bauer et al., 2013) and overlapping gene functions.

### SLAC1 is likely to play the role of a bicarbonate responsive protein

Our findings show that high intracellular bicarbonate could enhance SLAC1yc/OST1yn-mediated anion channel currents in *Xenopus* oocytes. These data indicated that either the OST1 protein kinase or SLAC1 or both proteins together may function as a bicarbonate responsive protein. To determine whether OST1 is essential for the 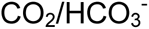 response, we co-expressed SLAC1 with the protein kinases CPK6 or CPK23 in *Xenopus* oocytes, as these protein kinases are well-known to activate SLAC1 in *Xenopus* oocytes (Geiger et al., 2010; Brandt et al., 2012). Four independent batches of oocytes showed that injection of high intracellular NaHCO_3_ or extracellular 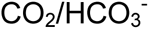 and co-expression of *β*CA4 and PIP2;1 with SLAC1 and CPK6 or CPK23 could enhance SLAC1 anion channel currents (Figures 8). Together these findings indicate that SLAC1 is likely to play the role of a bicarbonate responsive protein, but SLAC1 requires a protein kinase to mediate channel activity.

**Figure 8.**
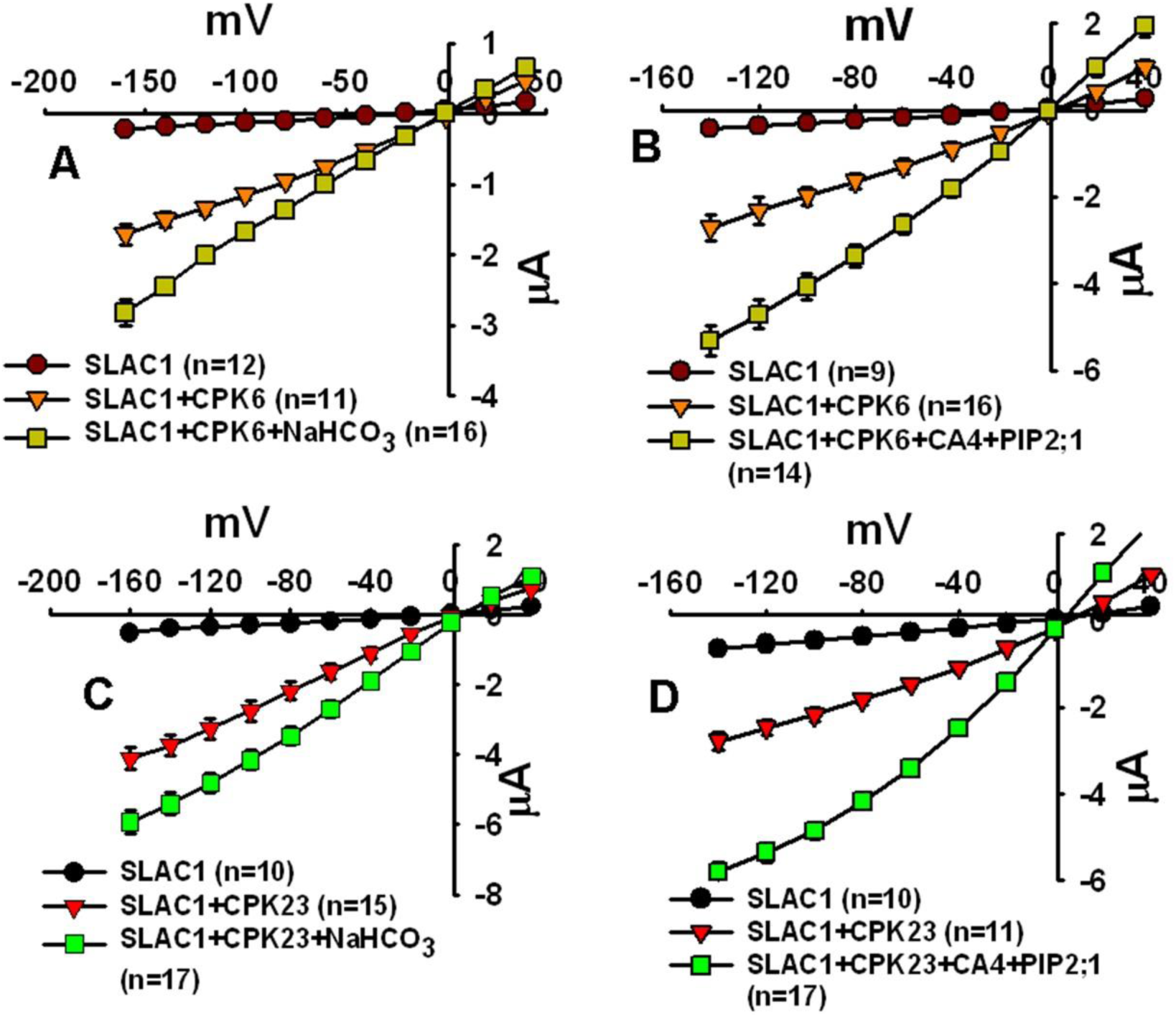
Intracellular bicarbonate enhances currents mediated by the SLAC1 anion channel in both CPK6-SLAC1- or CPK23-SLAC1-expressing oocytes and extracellular bicarbonate enhances currents in *β*CA4, PIP2;1 and CPK6-SLAC1or CPK23-SLAC1-co-expressing oocytes. **(A, C)** Bicarbonate injection into oocytes enhances SLAC1 anion channel currents when SLAC1 was co-expressed with the Ca^2+^-dependent protein kinases CPK6 or CPK23 rather than OST1 in oocytes. **(B, D)** *β*CA4 and PIP2;1 co-expression with 11.5 mM NaHCO_3_ in the bath solution enhances SLAC1/CPK6- and SLAC1/CPK23-mediated anion channel currents. Data are mean ± s.e.m. Results from four independent batches of oocytes showed similar results.

### RHC1 expression alone in oocytes produced ionic currents that were not affected by NaHCO_3_ or OST1

During the course of this research a recent study identified the MATE transporter RHC1 as a candidate 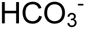 sensor when co-expressed with SLAC1, OST1 and HT1 in *Xenopus* oocytes (Tian et al., 2015). However, 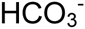 effects on oocytes expressing RHC1 alone or on oocytes expressing only OST1 and SLAC1 were not investigated. A full length RHC1 cDNA was obtained from a plant membrane protein cDNA collection (Jones et al., 2014). We then expressed the RHC1 cRNA in *Xenopus* oocytes with or without OST1. Unexpectedly, when expressing RHC1 alone oocytes showed ionic currents (Figure 9). The RHC1 MATE transporter-mediated currents were not affected by injection of 11.5 mM NaHCO_3_ into *Xenopus* oocytes in the presence or absence of OST1, and OST1 did not enhance RHC1-mediated currents (Figure 9).

**Figure 9.**
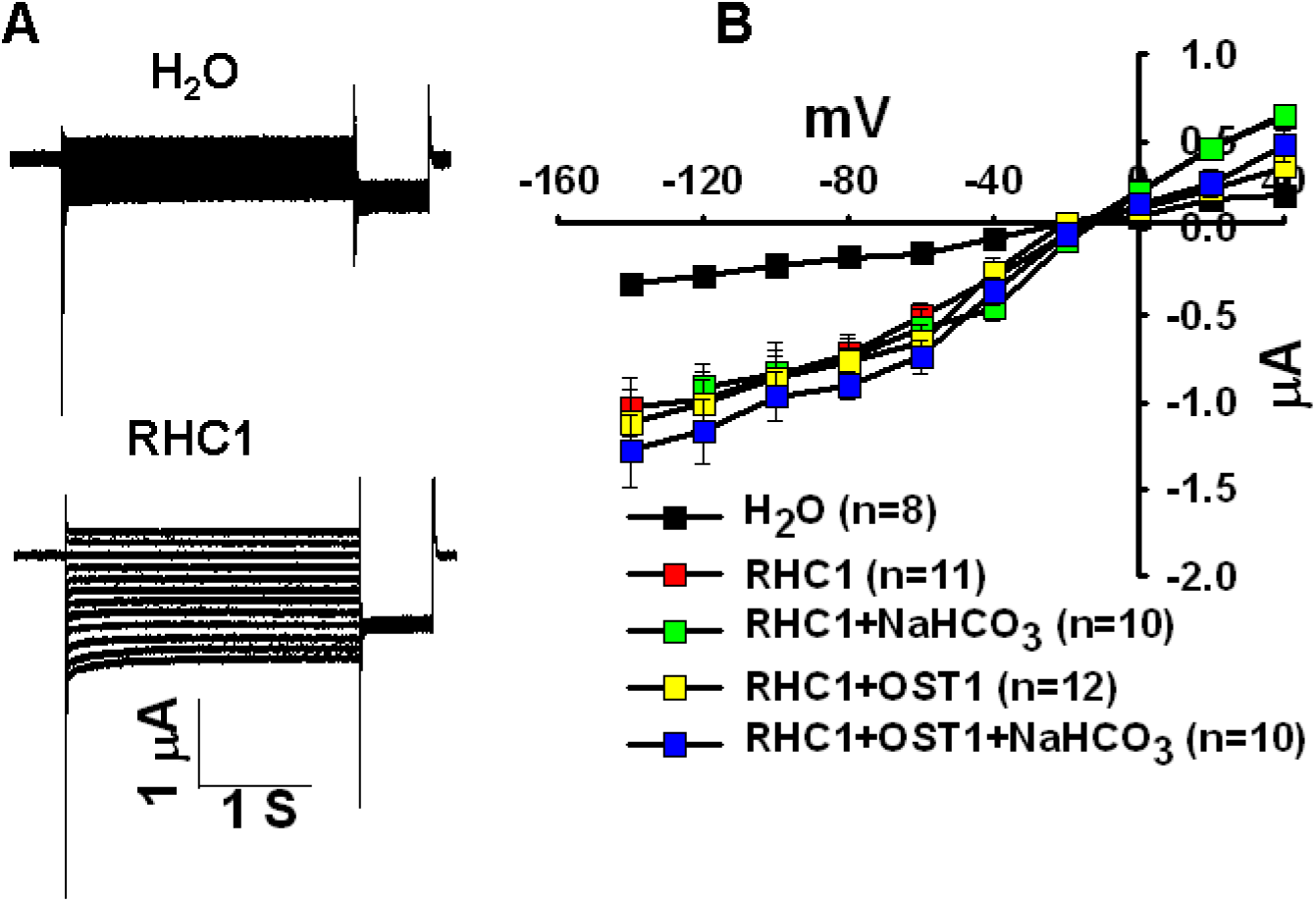
RHC1 expression in oocytes causes ionic currents and intracellular bicarbonate did not enhance RHC1-mediated anion channel currents in *Xenopus* oocytes in the presence or absence of the protein kinase OST1. **(A)** Whole-cell currents were recorded from oocytes expressing the indicated cRNAs. **(B)** Steady-state current-voltage relationships from oocytes recorded as in (A). Due to overlapping data of “RHC1” and “RHC1+NaHCO_3_” alternating data points are shown. Data are mean ± s.e.m. Results from three independent experiments showed similar results.

## DISCUSSION

The carbonic anhydrase *β*CA4 is involved in CO_2_-induced stomatal closing (Hu et al., 2010; Hu et al., 2015). To further investigate the CO_2_ signaling mechanisms mediated by the *β*CA4 protein, we found *β*CA4 interacted with PIP2;1.The interaction of PIP2;1 with *β*CA4 may explain why *β*CA4 has been found to be located at the intracellular side of the plasma membrane in plant cells even though *β*CA4 does not have a transmembrane domain (Fabre et al., 2007; Hu et al., 2015). It has been widely demonstrated that CO_2_ is transported across biomembranes via aquaporins: The tobacco aquaporin NtAQP1 displays CO 2 transport activity in *Xenopus* oocytes (Uehlein et al., 2003). The *Arabidopsis* AtPIP1;2 aquaporin displays CO_2_ permeability in a yeast expression system (Heckwolf et al., 2011). Four barley HvPIP2 aquaporins were recently shown to display CO_2_ permeability in *Xenopus* oocytes (Mori et al., 2014). Expression of PIP2;1 resulted in a change in the surface pH of oocytes (Musa-Aziz et al., 2009, 2014), showing that *Arabidopsis* PIP2;1 is permeable to CO_2_.

During the completion phase of the present study the RHC1 MATE transporter was identified and reported to function as a bicarbonate sensor candidate based on intracellular 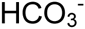 enhancement of SLAC1/OST1/RHC1/HT1-mediated anion channel currents in *Xenopus* oocytes (Tian et al., 2015). Whether SLAC1/OST1 co-expression alone also enables 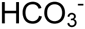-enhancement of SLAC1-mediated currents and whether RHC1 alone produces an electrogenic current was not investigated in this recent study. In the present study, we focused on the questions whether (1) high intracellular HCO_3_can enhance S-type anion channel currents in *Xenopus* oocytes? (2) Can we reconstitute extracellular 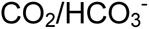 regulation of SLAC1-mediated anion currents? (3) Can we identify a candidate 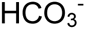 responsive protein?

Our experiments show that RHC1 expression alone mediates a clear ionic current in *Xenopus* oocytes that is not dependent on 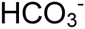 injection or OST1 co-expression. These findings are consistent with other plant MATE transporters that upon expression produce anion currents in *Xenopus* oocytes (Melo et al., 2013; Maron et al., 2010). Our experiments do not strictly exclude a role for RHC1 in CO_2_ signaling (Tian et al., 2015), but point to the need to investigate whether previous findings might result at least in part from additive currents mediated by SLAC1 and RHC1.

Intracellular bicarbonate enhances S-type anion channel currents in wild-type *Arabidopsis* guard cells (Hu et al., 2010; Xue et al., 2011; Tian et al., 2015). To mimic this process, we co-expressed SLAC1/OST1, SLAC1/CPK6 or SLAC1/CPK23 in *Xenopus* oocytes. We then microinjected NaHCO_3_ into oocytes at the same concentrations that enhance S-type anion channel currents in guard cells (Hu et al., 2010; Xue et al., 2011; Tian et al., 2015). We found that high intracellular 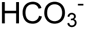could enhance SLAC1yc/OST1yn, SLAC1/CPK6 and SLAC1/CPK23-mediated anion channel currents in *Xenopus* oocytes. As the common protein in these analyses is SLAC1, these findings implicate SLAC1 as a candidate 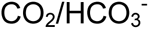 sensing protein.

The present findings show that enhancement of SLAC1 activity by intracellular 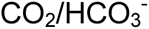 requires the presence of protein kinases (Figure 1). These findings are consistent with the requirements of protein kinase-mediated phosphorylation for activation of SLAC1 (Geiger et al., 2009; Lee et al., 2009; Geiger et al., 2010; Brandt et al., 2012; Hua et al., 2012) and the requirement of the OST1 protein kinase for CO_2_ signal transduction in plants (Xue et al., 2011; Merilo et al., 2013).

### More than one 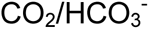 sensing pathway in guard cells

Note that the present findings do not exclude and indeed support the possibility of more than one 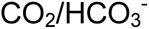 sensing mechanism in guard cells, as present and previous findings implicate the need for an active OST1 protein kinase in order for CO_2_ signal transduction to proceed (Xue et al., 2011; Merilo et al., 2013).

Furthermore, our present research shows that while NaHCO_3_ enhances the activity of SLAC1, significant SLAC1 activity prevails in the absence of NaHCO_3_ addition. Thus a second 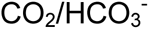 stimulated pathway would be required in guard cells that mediates activation of protein kinases that phosphorylate and activate SLAC1 (Geiger et al., 2009; Lee et al., 2009; Geiger et al., 2010; Brandt et al., 2012; Hua et al., 2012; Tian et al., 2015). This hypothesis is supported by a recent study and modeling together suggesting that two distinct CO_2_ signal transduction components exist in guard cells, one mediated by the plasma membrane located *β*CA4 and one dependent on the chloroplast-targeted *β*CA1 (Hu et al.,2015). Thus we propose that the direct 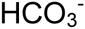-regulation of SLAC1 found here is not the only bicarbonate responsive protein in guard cells that contributes to CO_2_-induced stomatal closing.

### PIP2;1 and *β*CA4 are required for reconstitution of extracelluar 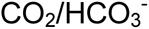regulation

Guard cell transcriptome studies have shown that multiple *PIP2* and *PIP1* aquaporin genes are expressed in guard cells under all investigated conditions (Leonhardt et al., 2004; Yang et al., 2008; Zhao et al., 2008; Bauer et al., 2013). These findings point to the hypothesis that higher order mutants in *PIP* aquaporin genes may be needed to affect stomatal CO_2_ responses. Further research will be needed to investigate this hypothesis.

When *β*CA4 and PIP2;1 together with SLAC1yc/OST1yn were co-expressed in *Xenopus* oocytes, SLAC1 anion channel currents were enhanced in the presence of extracellular NaHCO_3_. In contrast a non-functional PIP2;1 point mutant was identified here, PIP2;1-G103W. PIP2;1-G103W was targeted to the plasma membrane of oocytes, but did not enable extracellular NaHCO_3_-dependent enhancement of SLAC1 anion channel currents in *Xenopus* oocytes. The non-functional PIP2;1 point mutant PIP2;1-G103W has a mutation analogous to a mutation previously predicted to impair function of the grapevine VvPIP2;5 aquaporin (Shelden et al., 2009). Two models have been considered for the structural pathway by which CO_2_ is transported by aquaporins: (1) CO_2_ may be transported via a central pore formed by an aquaporin tetramer. (2) An alternative model has been considered in which CO_2_ is transported via the same channel pore as water in aquaporins (Wang et al., 2007; Horsefield et al., 2009). In the present study, the *PIP2;1G103W* mutation disrupted both water and CO_2_ transport, which might be interpreted to support a common pore model for water and CO_2_ transport. However, more in depth studies would be needed to carefully investigate these two models, as one mutation is insufficient to make a strong conclusion.

The extracellular 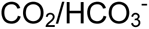 enhancement of SLAC1yc/OST1yn-mediated anion channel currents by *β*CA4 and PIP2;1 requires functional PIP2;1. We thus were able to reconstitute extracellular 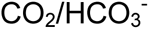 regulation of SLAC1-mediated anion currents in *Xenopus* oocytes by co-expression of *β*CA4, PIP2;1, SLAC1 and either OST1, CPK6 or CPK23. We propose a working model for mechanisms contributing to the CO_2_ signalling pathway leading to stomatal closure. When the CO_2_ concentration in leaves (*C*i) is elevated, CO_2_ influx across the plasma membrane of guard cells is enhanced through aquaporins. The carbonic anhydrases accelerate the production of intracellular 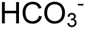, and elevated intracellular 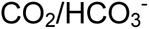 enhances SLAC1 anion channel activity, triggering the closure of stomatal pores. When *Ci* is low, as occurs during the light phase (Hanstein et al., 2001), the intracellular 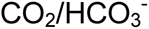 concentration is reduced and S-type anion channel activity is reduced.

## METHODS

### Two-electrode voltage-clamp recordings in *Xenopus* oocytes

All constructs were cloned into the pNB1 oocyte expression vector using the USER method (Nour-Eldin et al., 2006). cRNAs were synthesized from 0.5-1 μg of linearized plasmid DNA template using the mMessage mMachine *in vitro* Transcription Kit (Ambion). Approximately 20 ng of each indicated cRNA was injected into oocytes for voltage-clamp recordings. Injected oocytes were incubated in ND96 buffer at 16°C for 2-3 days prior to electrophysiological recording. The ND96 buffer contained 10 mM MES/Tris (pH 7.5), 1 mM CaCl_2_, 1 mM MgCl_2_ and 96 mM NaCl. Whole-cell ionic currents were recordings with a Cornerstone (Dagan) TEV-200 two-electrode voltage clamp amplifier and digitized using an Axon Instruments Digidata 1440A Low-Noise Data Acquisition System (Molecular Devices) controlled by pClamp acquisition software (Molecular Devices Sunnyvale, CA, USA). Microelectrodes were fabricated with a P-87 Flaming/Brown microelectrode micropipette puller (Sutter, Novato, CA) from borosilicate glass (GC200TF-10; Warner Instruments, Hamden, CT, USA) and the tips were filled with 3M KCl. The resistance of the filled electrodes was 0.5–1.5 m Ω.

As oocytes batches vary in the protein expression level among batches and thus the magnitude of ionic currents from one week to another varies, the indicated controls were included in each batch of oocytes and control conditions were recorded intermittently with investigated conditions to avoid a time of measurement dependence of data. Furthermore, data from one representative batch of oocytes are shown with controls in each figure, and were reproduced in at least three independent oocyte batches as indicated in results. The numbers of oocytes recorded for each condition in the depicted batch of oocytes are also provided in the figures. Oocytes were recorded in 10 mM MES/Tris (pH 7.4), 1 mM MgCl_2_, 1 mM CaCl_2_, 2 mM KCl, 24 mM NaCl, and 70 mM Na-gluconate buffer. Osmolality was adjusted to 220 mM using D-sorbitol. Steady state currents were recorded starting from a holding potential of 0 mV and ranging from +40 to -160 mV in -20 mV decrements, followed by a −120 mV voltage “tail” pulse. Note that the time-dependent properties of SLAC1 channel in *Xenopus* oocytes vary among individual oocytes. This variability in time-dependent properties was noted in early studies of S-type anion channel currents in guard cells (Schmidt and Schroeder, 1994), and may depend on posttranslational modification of the channel protein that requires further analyses. Data were low-pass-filtered at 20 kHz throughout all recordings. For application of intracellular bicarbonate, NaHCO_3_ was injected into each oocyte and the final concentration is given in the figures caption. The concentrations of free bicarbonate and CO_2_ were calculated using the Henderson–Hasselbalch equation 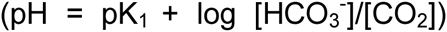, where 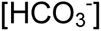 represents the free bicarbonate concentration and [CO_2_] represents the free CO_2_ concentration; pK_1_ = 6.352 was used for the calculation (Speight, 2005). Injection of 11.5 mM NaHCO_3_ results in 10.5 mM free 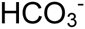 and 1mM free CO_2_ at pH 7.4. The calculation of intracellular bicarbonate was based on cytosolic oocyte volumes ≈ 500 nL, and the volume of injected NaHCO_3_ buffer was 50 nL. Bicarbonate microinjections were performed 20 min before voltage-clamp experiments. All experiments were performed at room temperature. Surface pH measurements and the water swelling assay in *Xenopus* oocytes were performed as described in (Geyer et al., 2013; Musa-Asziz et al., 2014). Briefly, a baked, silanized (*bis*-di-(methylamino)-dimethylsilane; Sigma-Aldrich, cat. no. 14755) borosilicate glass microelectrode, with a H^+^ Ionophore I, mixture B, (Sigma-Aldrich, cat. no. 95293) liquid membrane at its 15 μm (inner diameter) tip, and backfilled with a solution (containing, in mM, 40 KH_2_PO_4_, 23 NaOH, 15 NaCl, adjusted to pH 7.0) was used to measure pH_s_. The pH_s_ microelectrode was connected to a FD223 electrometer; (World Precision Instruments, Sarasota, FL, USA) and mounted on an ultrafine micromanipulator (model MPC-200 system; Sutter Instrument Co., Novato, CA) to position the pH_s_-electrode tip at the surface of the oocyte. To record pH_s_, the tip was then advanced a further ~40 μm, creating a visible dimple on the oocyte membrane. Periodically the electrode tip was withdrawn ~300 μm for recalibration in the bulk extracellular fluid (pH 7.50). Composition of ND96 and 5% CO_2_/33 mM 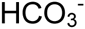 solutions were as described in Musa-Aziz et al. (2014) and flowed into the chamber at 4 ml/min.

### Confocal Microscopy

Confocal imaging of PIP2;1-YFP fusion proteins expressed in *Xenopus* oocytes was acquired by spinning-disc confocal microscopy (Nikon Eclipse TE2000-U). Images show an optical slice of each oocyte. Fluorescence imaging Images were captured with an electron multiplying charge-coupled device (EMCCD) camera (Cascade II: 512, Photometrics, Tucson, AZ, USA) using Metamorph software (Universal Imaging, Downington, PA, USA).

### Yeast two-hybrid Screen

For isolation of *β*CA4-interacting proteins, a yeast two-hybrid screen was conducted. *βCA4* was fused with GAL4-BD in the vector pGBKT7 (Causier and Davies, 2002). pGBKT7-*βCA4* was used as bait to screen for interacting proteins from a normalized commercial *Arabidopsis* cDNA library constructed in the pGADT7 vector (Clontech). pGBKT7-*βCA4* was transformed into the yeast strain Y2HGold and the library was in the Y187 strain. The mating method was adopted and diploid cells were grown on the SD-Leu-Trp medium plus 40 mg/mL x-gal at pH 5.8. Interactions were detected as blue cells, but no robust yeast 2-hybrid interactors of *βCA4* were identified in this screen.

### Split-ubiquitin system analyses

The split-ubiquitin system was adopted and improved to test direct protein-protein interactions and to screen *β*CA-interacting proteins from an *Arabdopsis* cDNA library (Grefen et al., 2007). The *β*CA4 cDNA was cloned into the vector pMetYC-DEST. pMetYC-*β*CA4 was used as a bait to screen an *Arabdopsis* cDNA library, which was constructed in the prey vector pNX33-DEST. The THYAP4 yeast colonies transformed with pMetYC-*β*CA4 were mated with the THYAP5 yeast transformed with 3 μg of *Arabidopsis* cDNA library (Grefen et al., 2007). The co-transformants were grown on SD-Leu-Trp medium plus 40mg/mL x-gal at pH5.8. Blue colonies indicated putative interactions.

### BiFC experiments in *N. benthamiana*

For split YFP complementation assays (BIFC), the vectors pXCSG-YN155 and pXCSG-YC84 were used (Chen et al., 2008). To generate the *βCA4*-YC84 and *PIP2;1*-YN155 constructs, full-length *PIP2;1* cDNA without stop codons was amplified and cloned into the binary vector pXCSG-YN155, and a 836 bp *βCA4* cDNA without stop codons was amplified and cloned as a fusion with the binary vector pXCSG-YC84. Split luciferase assays were carried out as described (Chen et al., 2008). Briefly, the *PIP2;1* cDNA was cloned into a vector containing the C-terminal half of luciferase (cLUC) and *βCA4* was cloned into the N-terminal half of luciferase (nLUC). These constructs were transformed into the *Agrobacterium* strain GV3101 and then co-infiltrated into *N. benthamiana* leaves with P19 at an OD_600_ of 0.8. After three days of infiltration, the infiltrated leaves were harvested for bioluminescence detection. Images were captured with a CCD camera.

### Co-immunoprecipitation experiments in *N. benthamiana*

The *Agrobacterium* strain GV3101 with the helper plasmid pMP90K carrying *β*CA4 and PIP2;1 was co-infiltrated at an OD_600_ of 0.8 together with the p19 strain in *N. benthamiana*. Protein extraction was performed with infiltrated leaves after three days of infiltration (Nishimura et al., 2010). Infiltrated leaves were then sprayed with water containing 0.01% Silwet L-77 after 3 days infiltration before leaf excision. 1.0 g leaves were ground in liquid nitrogen and the powder tissues were re-suspended with 2.0 mL of extraction buffer (50 mM Na-phosphate (pH7.4), 1 mM DTT, 0.1% NP-40,150 mM NaCl, and 1×protease inhibitor cocktail). Crude extracts were centrifuged at 18,000 g for 20 min at 4°C. Supernatant passed through Miracloth (Calbiochem) was used for immunoprecipitation as an input. Input proteins were incubated with 50 μL anti-HA matrix for 3 h at 4°C and the immune-complexes were washed four times with 500 μL washing buffer (50 mM Na-phosphate (pH7.4), 150 mM NaCl, 0.1% NP-40). Proteins were separated by 10% SDS-PAGE gel and electrotransferred onto immobilon-P membrane. Membrane was blocked overnight in PBS-T buffer with 5% skim milk (Biorad), then washed three times with PBS-T buffer and incubated with anti-GFP or anti-HA antibodies. The membranes were incubated with 1:10000 diluted anti-mouse HRP-conjugated for 1 h. Bio-Rad’s Clarity ECL western blotting substrate was then used to perform detection.

### Time-resolved intact leaf stomatal conductance experiments with [CO_2_] shifts

To identify the *PIP2;1* T-DNA insertion mutation, we used two qPCR primers: F: TACCACCAATTCGTTCTG; R: AGAGATCACAACTTCATTTATTC. Four to six week old plants growing in a growth chamber at 70% humidity were used for intact-leaf CO_2_-induced stomatal conductance change analyses as previously described (Hu et al., 2010). Briefly, stomatal conductance was firstly stabilized at 400 ppm of [CO_2_] and recorded for an additional 30 min, then [CO_2_] was shifted to elevated CO_2_ for 30 min and then again changed to 100 ppm. The data shown are means of 4 leaves per genotype per treatment ± s.e.m. Relative stomatal conductances were calculated by normalization with respect to the average of the final ten data points at 400 ppm [CO_2_] before elevating [CO_2_].

### Stomatal aperture measurements

Three week-old *Arabidopsis* plants grown in a growth chamber at 70% humidity were used for analyses of stomatal movements in response to ABA. Intact leaf epidermal layers with no mesophyll cells in the vicinity were prepared as described (Hu et al., 2010; Xue et al., 2011). Leaf epidermal layers were preincubated for 2 h in opening buffer (10 mM MES, 10 mM KCl, 50 μM CaCl_2_ at pH 6.15) and then incubated with buffers supplemented with 10 μM ABA. For time course analyses, individual stomata were imaged and individually tracked at different time points. Stomatal apertures were measured using ImageJ (Schneider et al., 2012). Data shown are from genotype blind analyses (n = 3 experiments, 30 stomata per experiment and condition).

### Simulation of CO_2_ transport in oocytes

Our model describes the dynamics of CO_2_ influx using a set of differential equations and is detailed in previous work (Somersalo et al., 2012). Briefly, it consists of a spherical oocyte of radius 650 μm surrounded by a layer of unconvected extracellular fluid of thickness d=100 μm in which reactions can take place. The oocyte and fluid layer are immersed in a bath in which the concentration of the reactants are assumed to be constant. Within the layer and the oocyte the model describes the formation and disassociation of carbonic acid as:

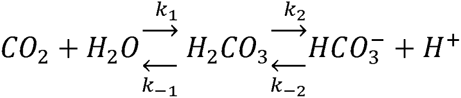

where *k_1_, k_-1_, k_2_, and k_-2_* are rate constants. In addition, we assumed that there is one non 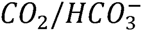 buffer, denoted by *HA/A^−^*.

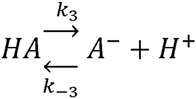

We assumed that the membrane is permeable to CO_2_ and incorporated intracellular carbonic anhydrase-like activity by multiplying the rate constants *k_1_* and *k_-1_* by an acceleration factor A. Assuming spherical symmetry, the rate equations corresponding to the above reactions can be solved along a radial line using the methods-of-line algorithm. All parameters and initial conditions were chosen as in the study by *Somersalo et al.,* 2012 with the exception of the membrane permeability P_M CO2_, which was chosen to be lower.

## SUPPLEMENTAL FIGURE

**Supplemental figure 1.** Injection of 11.5 mM NaHCO_3_ (pH 7 or pH 8) into oocytes also causes enhancement of SLAC1-mediated anion channel currents, whereas 23 mM sorbitol has no effect on SLAC1 activity. Data are mean ± s.e.m.

**Supplemental figure 2.** Steady-state current-voltage relationships show the average magnitude of SLAC1yc/OST1yn-mediated anion channel currents recorded from oocytes injected with 11.5 mM NaCl were reduced rather than enhanced. Data are mean ± s.e.m. Data are representative of experiments performed on three independent oocyte batches.

**Supplemental figure 3.** Surface pH (pH_s_) measurements from oocytes exposed to 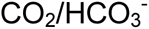. Oocytes were exposed to 5% CO_2_/33 mM 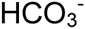 long enough for the pH_s_ to rise and then decay exponentially to a stable value. Traces from oocytes recorded in the same batch are shown.

**Supplemental figure 4.** The *PIP2;1-W85A and PIP2;1-F210A* mutation isoforms did not impair the PIP2;1-CA4 enhancement of SLAC1/OST1-mediated anion channel currents in oocytes by extracellular 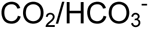. **(A)** Whole-cell currents were recorded from oocytes expressing the indicated cRNAs with 11.5 mM NaHCO_3_ in the bath solution. The voltage protocol was the same as in Figure 1. **(B)** Steady-state current-voltage relationships from oocytes recorded as in (A). Data are mean ± s.e.m. Results from three independent batches of oocytes showed similar results.

**Supplemental figure 5** Simulated membrane surface pH (pH_s_) as a function of time for baseline parameter values (black curve), in the presence of intracellular carbonic anhydrase (CA) activity (red curve), for simulated increased membrane CO_2_ permeability (green curve), and in the presence of both intracellular CA activity and increased membrane CO_2_ permeability (blue curve).

**Supplemental figure 6 (A)** Structure of *PIP2;1* gene and T-DNA insertion. *PIP2;1 c*onsists of four exons (open boxes); black boxes highlight the 5’ and 3’ untranslated regions, respectively. Mutant line *pip2;1* (ABRC stock name CS320492) has a T-DNA insertion in 5’-UTR region. **(B)** qPCR analyses showed that *pip2;1* was a knockdown mutant. Expression level was compared to EF-1a.

## ACKNOWLEDGEMENTS

We thank Aaron Stephan (University of California San Diego), Benjamin Brandt (University of Geneva, CH) and Fraser Moss (Case Western Reserve University) for comments on the manuscript. This research was funded by a grant from the National Science Foundation (MCB1414339) to J.I.S and W.J.R, and in part supported by grants from the National Institutes of Health (GM060396-ES010337) to J.I.S. and the China National Natural Science Foundation (31271515) and the Fundamental Research Funds for the Central Universities (2662015PY179) to H.H. Surface pH experiments were supported by American Heart Association Postdoctoral Fellowships (AHA09POST2060873 and AHA11POST7670014) to X.Q. and Office of Naval Research Grants (N00014-11-1-0889; N00014-14-1-0716; N00014-15-1-2060), the National Institutes of Health (U01-GM111251) and the Meyer/Scarpa Chair to W.F.B.

## AUTHOR CONTRIBUTIONS

C.W performed all of the oocyte TEVC electrophysiological experiments and data analyses. H.H. preformed S.U.S., bimolecular fluorescence complementation and co-immunoprecipitation experiments at UCSD. X.Q. and B.Z. performed surface pH experiments and C.W., X.Q. and B.Z. performed Pf measurements on *Xenopus* oocytes. D.X. performed split luciferase experiments. W.J.R. constructed the mathematical model. The project was conceived by J.I.S. and C.W., H.H., W.F.B., and J.I.S. contributed to the research design, provided support and suggestions throughout the research and manuscript preparation. C.W. and J.I.S. wrote the manuscript with contributions from other authors.

